# A Predictive Model: The Key to Success for Badminton Servers and Receivers

**DOI:** 10.1101/2024.02.10.579788

**Authors:** Cheng Liu

## Abstract

To explore the influence of technical actions on badminton competition outcomes, this study analyzes the frequency and impact of specific movements made by players, distinguishing between the server and receiver roles. Focusing on international competitions from 2019 to 2023, we collected data on 23 distinct technical actions (e.g., Net Front, Slice/Drop, Push) to construct a predictive model. The study distinguishes itself by employing a Random Forest algorithm to ascertain the significance of each technical action, utilizing forward stepwise selection and 5-fold cross-validation for feature refinement. SHAP value analysis further validated the pivotal roles of ‘Net Front’, ‘Slice/Drop’, and ‘Push’ across both sexes, linking higher frequencies of ‘Net Front’ with increased match-winning probabilities. Model validation on a test set demonstrated effective performance in both sexes, with the model based on male data exhibiting higher accuracy and predictive values, surpassing the performance of the female data model. This comprehensive examination, grounded in quantitative analysis, not only enhances our understanding of badminton gameplay dynamics but also offers valuable insights for coaching strategies and training methodologies.

## INTRODUCTION

In badminton matches, the adoption of appropriate game strategies plays a decisive role in securing victory. Particularly, the first serve of each game significantly affects the pace and control of the match. The serving player’s strategic choice not only has a profound impact on the initial rhythm of the game but also largely determines the control over the match. The technical actions of the server directly reflect their tactical style and have an immediate influence on the early direction of the game. Likewise, the responding strategy of the receiver is equally critical. How the server effectively utilizes their serving advantage, as well as how the receiver adjusts their strategy to counter the server’s technical actions, are key factors that determine the match rhythm and outcome. However, current research lacks a quantitative analysis of the differences in technical action frequencies between the server and receiver, and their specific impacts on match outcomes, limiting our ability to deeply understand and analyze badminton match strategies.

This study aims to bridge this gap by thoroughly analyzing the differences in technical action frequencies between servers and receivers in each game, exploring their impacts on match outcomes to construct a model capable of predicting match results. We will focus on collecting and analyzing relevant data from high-level badminton matches, paying particular attention to the technical action frequencies during the first serve and reception in each game, with the goal of uncovering the relationship between these actions and match outcomes. By identifying the key technical actions that decisively influence match results, this study not only provides a scientific basis for coaches and players to adjust strategies during matches but also hopes to contribute new theoretical and practical insights into the field of badminton training and match strategy research.

## Methods

### Data collection

This study selected videos of men’s and women’s singles badminton matches above the Super750 level from 2019 to 2023 for observation and analysis of players’ technical and tactical levels, encompassing major international badminton events such as the 2020 Tokyo Olympics, World Championships, All England Championships, Sudirman Cup, Thomas Cup, and Uber Cup. A total of 153 matches comprising 358 games for men and 150 matches comprising 344 games for women were analyzed. The selected match videos primarily came from tracking national team competitions, as well as from sources such as Bilibili, CCTV Video, and the official website of the Badminton World Federation (BWF).

The study employed video observation methods to statistically analyze technical actions and match outcomes. Four national second-level badminton players from our research team served as data recorders. Initially, the collected videos were uploaded to the APP of ‘match-analytic’ badminton technique and tactics statistical analysis software, developed in collaboration with Shanghai Danzhu Sports Technology and Culture Co., Ltd. Utilizing this software. Subsequently, the four staff members independently observed and recorded statistics from the videos. The results were then cross-verified. In cases of inconsistency, the four staff members would review together until a consensus was reached. This integrated approach, leveraging both manual observations and sophisticated software analysis, ensured the accuracy and reliability of the data collected, providing a solid foundation for analyzing the technical and tactical levels of players in high-level badminton competitions.

### Preprocessing

After collecting the data, we first conducted a statistical analysis of the types of technical actions in all matches, identifying a total of 23 initial features (High, Smash, Dribble, Push, Slice/Drop, Lift, Block Smash, Net Front, Clear, Net Drop from Lift, Lift from Slice, Pull, Hook, Slice Lift, Block, Lift Smash, Block Hook, Hook from Lift, Drive, Net Drop from Slice Drive, Lift from Slice Drive, Flat High, and Block from Slice Drive). Then, we calculated the frequency of technical actions for both the serving side and the receiving side in each game to construct vectors. A feature vector was formed by subtracting the vector of the receiving side from that of the serving side. Furthermore, we defined the labels based on the outcome of each game: if the serving side won the game, the label was set to 1; if the receiving side won, the label was set to 0.

### Model description

We implemented a random forest classifier using the scikit-learn package in Python 3.97 programming language (https://www.python.org/). To generate the training (70%) and test (30%) datasets, we split the dataset using the train_test_split function, which randomly partitions a dataset into training and test subsets with test_size=0.3 parameter. We applied this function to each sex individually. To identify the key technical actions critical for predicting match outcomes, we first constructed a random forest classifier using the default parameters based on the entire training dataset. We extracted the feature importance parameters from this classifier and sorted the features according to their importance. Subsequently, the features were incrementally included in the default random forest classifier based on their ranked importance. This process was validated using a 5-fold cross-validation method to determine the optimal number of technical actions for describing the model features. To build a random forest classifier with the best hyperparameters, we implemented the exhaustive grid search approach using the GridSearchCV function to the training dataset with five-fold cross-validation. A total of 10,000 random forest classifier models were evaluated with different combinations of hyperparameters: max_features="auto"; n_estimators ranging from 100 to 1,000 with an interval of 100; max_depth ranging from 2 to 20 with an interval of 2; min_samples_leaf ranging from 2 to 20 with an interval of 2; and min_samples_split ranging from 2 to 20 with an interval of 2. As a result, we selected the male model with n_estimators=600, max_depth=8, min_samples_leaf=2 and min_samples_split=2 hyperparameters, which showed the highest average AUC at 0.9656. Also, the female model was built with n_estimators=1000, max_depth=12, min_samples_leaf=2 and min_samples_split=8 hyperparameters, which showed the highest average AUC at 0.8950.

## RESULTS

### Identification of Critical Technical Movements Contributing to Success in Competitions

In this study, we collected data from international badminton competitions from 2019 to 2023, aiming to statistically analyze the frequency of each technical action in each rally of the competitions. The considered technical actions include High, Smash, Dribble, etc, totaling 23, see Methods for details. The frequency of these technical actions was used as feature vectors to analyze the winning patterns of both the serving and receiving sides, further aiming to build a model capable of predicting competition outcomes.

The data, divided by sex, years, and competition outcome, was split into a training set and a test set with a ratio of 7:3. Specifically, the training set for men’s competitions contained 250 samples, while the test set contained 108 samples; the women’s competitions training set included 240 samples, and the test set contained 104 samples. This data division method ensured sufficient model training and reliable test results (Fig 1).

**Figure 1:**
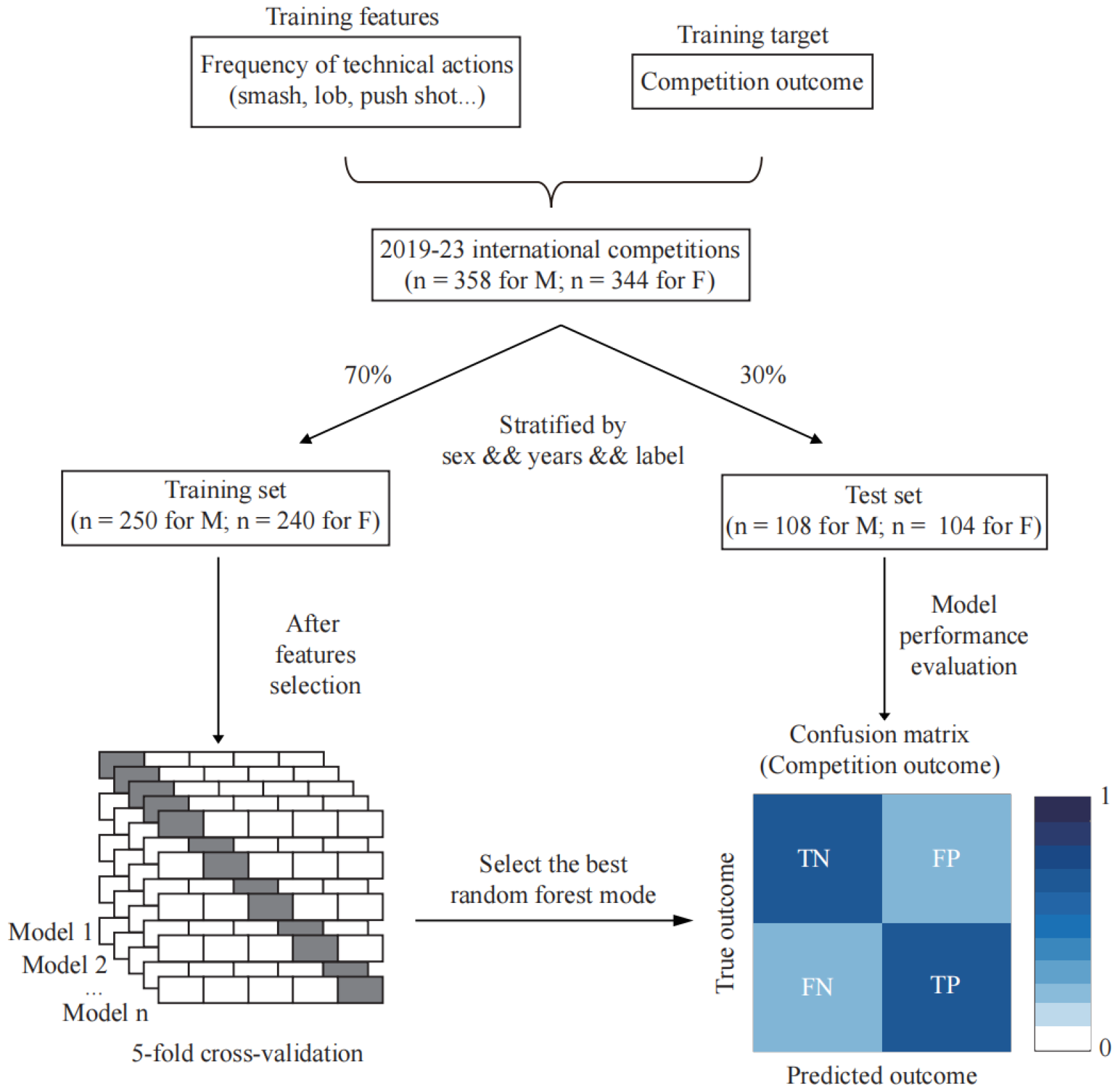
overview of study.

For the first time in this study, a Random Forest algorithm was used on the training set to build the model, evaluating the importance of each technical action feature. Through these evaluation results, we ranked the features by importance and used forward stepwise selection combined with 5-fold cross-validation to filter out key features that significantly impact the model’s performance. Using the AUC (Area Under the Curve) as the performance evaluation metric, it was found that for male athletes’ data, the model’s performance reached its optimum upon introducing the most critical five features (Net Front, Slice/Drop, Flat High, High, Push), with an AUC value of 0.9726. In contrast, for female athletes’ data, the model’s AUC value reached 0.8730 after introducing the top 22 most important features (Net Front, Slice/Drop, Flat High, High, Push, Smash, Block Smash, Drive, Block, Hook, Lift Smash, Lift, Dribble, Net Drop from Lift, Pull, Block Hook, Hook from Lift, Slice Lift, Lift from Slice, Lift from Slice Drive, Clear, Net Drop from Slice Drive) (Fig 2A-B). This result reflects the tactical styles of female badminton players in serving and receiving are more complex compared to their male counterparts to a certain extent. Finally, we built the final model after determining the optimal hyperparameters based on 5-fold cross-validation.

**Figure 2:**
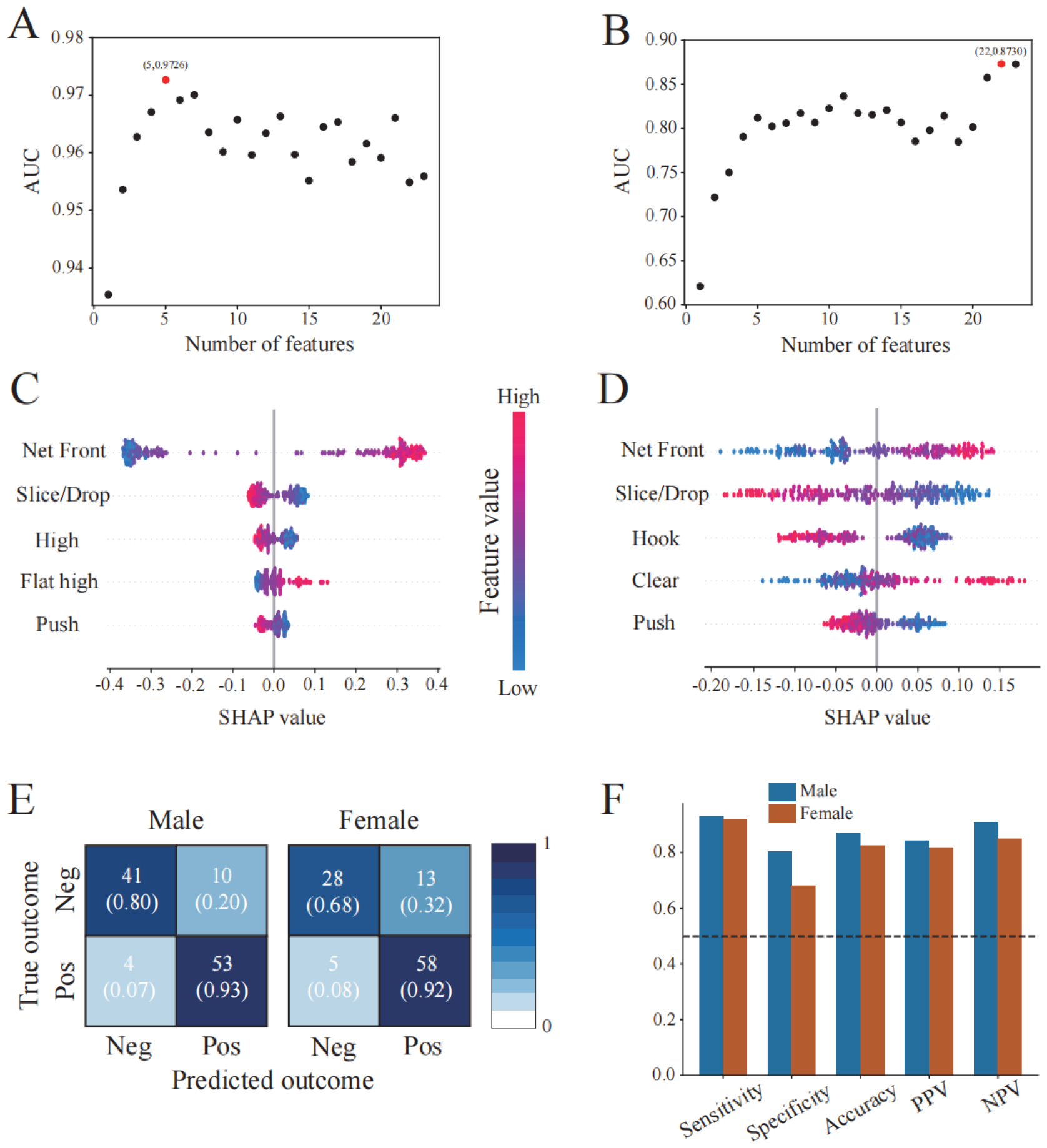
feature selection and model evaluation. (A-B) Effect of Incremental Feature Addition on male/female model AUC, respectively. (C-D) SHAP values of top 5 important features on male/female model, respectively. (E) Confusion matrix for models. (F) Model evaluation performance illustrated by sensitivity, specificity, accuracy, PPV and NPV.

### Quantitative Analysis and Model Performance Validation of the Impact of Key Technical Actions on Badminton Competition Outcomes

After the model was constructed, to further understand the specific impact of each feature on the outcomes of competitions, we calculated the SHAP values for each feature. The analysis showed that among the top five key features evaluated, the technical actions of ‘Net Front’, ‘Slice/Drop’, and ‘Push’ had a significant impact on the outcomes of both men’s and women’s competitions, presenting a consistent pattern. Particularly for ‘Net Front’, its SHAP values increased with the value, indicating that a higher frequency of this technical action by the serving side compared to the receiving side is more likely to lead to victory in the competition. Conversely, the situations for ‘Slice/Drop’ and ‘Push’ were opposite (Fig 2C-D).

When evaluating the model’s performance based on the preserved test set, the following results were obtained: for males, Sensitivity was 0.9298, Specificity was 0.8039, Accuracy was 0.8704, Positive Predictive Value (PPV) was 0.8413, and Negative Predictive Value (NPV) was 0.9111; for females, Sensitivity was 0.9206, Specificity was 0.6829, Accuracy was 0.8269, PPV was 0.8169, and NPV was 0.8485. These results indicate that the model captures the technical action features of male players more accurately compared to females, consistent with the findings from the feature selection phase, and both significantly exceeded the baseline of 0.5 (Fig 2E-F).

### Future Plans

1.To visualize and apply the results of this study on a web platform, enabling future researchers and coaches to input required features in designated input fields. The website will provide specialized feedback based on the outcomes of this research, facilitating the adjustment of tactical strategies before competitions.

## DISCUSSION

This study initiates from the perspective of the serving and receiving players’ technical actions in each game, elucidating the respective key influencing factors of both parties in the competition process and constructing a predictive model based on these insights for subsequent researchers and coaches to study and reference. By meticulously analyzing the frequency of 23 distinct technical actions in international badminton competitions and employing a sophisticated Random Forest algorithm, we have not only identified the pivotal technical actions that significantly impact match outcomes but have also quantified their influence through the calculation of SHAP values. This innovative approach has allowed us to reveal that technical actions such as ‘Net Front’, ‘Slice/Drop’, and ‘Push’ play a crucial role in determining the victory, offering a nuanced understanding that transcends traditional analyses. The competition prediction models were also constructed measured by Sensitivity, Specificity, Accuracy, PPV and NPV, which could be directed applied by future related researchers and athletes.

This study delves into the winning patterns of serving and receiving players in badminton matches by analyzing the frequency of technical actions, highlighting the significance of key technical maneuvers in determining match outcomes. However, the factors influencing the results of badminton matches extend far beyond these, including but not limited to the athletes’ physical fitness, psychological state, tactical layout, opponents’ playing styles, and the environment of the competition. For instance, an athlete’s physical condition is crucial for endurance and speed on the court, while psychological factors like confidence and stress significantly impact performance. Furthermore, strategic tactical planning and choice of strategy can effectively counter various opponents, optimizing match performance.

Future research should build upon the findings of this study to further explore and analyze how these non-technical factors influence match outcomes. Such investigations can provide athletes with more comprehensive training and competition strategies and allow coaches to understand the keys to winning strategies more deeply, enabling more precise formulation of tactics during training and competitions to improve win rates. By considering technical actions along with the aforementioned multitude of factors, we can gain a more comprehensive understanding of the key determinants of victory in badminton matches, contributing new theoretical and practical insights to the development of the sport.

